# Shear-Induced Oscillations and Hydrodynamic Buffering Stabilize Sperm Surface Navigation

**DOI:** 10.64898/2026.06.03.729852

**Authors:** Mengna Liu, Antai Tao, Rongjing Zhang, Junhua Yuan

## Abstract

Navigation near boundaries under strong flow is central to microswimmer transport in active matter. Using microfluidics, we track rolling bovine sperm near planar walls in Poiseuille flow with near-wall shear rates up to 50 s^−1^. As flow increases, we observe a universal dynamical transition: circular surface swimming at zero flow, upstream rheotaxis at weak flow, and a novel near-surface oscillation (NSO) state at high shear, characterized by large-angle oscillations and periodic lifting from the wall. Surprisingly, sperm remain concentrated in a near-wall layer even when downstream advection dominates. A minimal mechanistic model combining hydrodynamic wall interactions, shear-driven Jeffery rotation, and steric flagellar-wall collisions reproduces these transitions and reveals a hydrodynamic buffer zone—a range of shear rates where the mean wall distance remains nearly constant. This buffering arises from a competition between wall-attracted and bulk-oscillatory states, providing a robust physical mechanism for surface navigation in fluctuating environments.

## Introduction

The transport of active particles in external fields is a central theme in non-equilibrium physics, with deep implications for biological function. Sperm motility represents a paradigmatic system, where cells must navigate the complex, dynamic environment of the female reproductive tract to fertilize an egg [1-4]. While mechanisms such as chemotaxis [5] and thermotaxis [6] contribute, rheotaxis—the ability to orient and swim against a flow—is considered the dominant mechanism for long-range guidance [7-10]. Previous work has established that sperm exhibit rheotaxis near surfaces under weak flows (shear rate < 10 s^−1^) [11-13]. However, physiological flows are often strong and highly variable, driven by muscle contractions and intricate topography [9,14-17]. How sperm maintain upstream progress and surface proximity in high-shear regimes remains an open question.

Microorganisms near boundaries typically exhibit surface accumulation [18-21] and circular trajectories [12,22], behaviors governed by far-field hydrodynamic interactions (force and rotlet dipoles) [23]. Under flow, these interactions compete with shear-induced rotation, often described by Jeffery orbits [24-28]. While bacteria in strong flows display oscillatory trajectories near surfaces due to chirality and shape [29], sperm possess distinct flagellar mechanics and beating patterns [30-35]. The interplay between flagellar elasticity, near-wall hydrodynamics, and high-shear flow creates a rich, unexplored phase space of motility. Although simulations suggest diverse behaviors under varying flow conditions [36-39], the 3D dynamics of sperm in high-intensity flows remain experimentally uncharacterized.

Here, we investigate the dynamics of bovine sperm in planar Poiseuille flows with near-wall shear rates up to 50 s^−1^. We identify a universal transition sequence: from circular swimming, to stable rheotaxis, and finally to a novel near-surface oscillation (NSO) mode. Surprisingly, we find that sperm resist hydrodynamic washout by maintaining a stable mean distance from the wall across a broad range of high shear rates. We explain this via a hydrodynamic buffering mechanism, validated by a mechanistic model integrating Jeffery orbits, wall interactions, and steric flagellar collisions. This mechanism stabilizes surface navigation in fluctuating environments, offering a robust physical framework for understanding microswimmer transport in complex flows.

### Flow-dependent Transitions in Near-surface Sperm Motility

We generated planar Poiseuille flow in a microfluidic chamber using a syringe pump (Fig. 1a and Fig. S1a). Calibration of the mean bulk flow speed 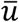 versus pumping rate *q* (Fig. S1b) indicates that, in our experiments, the maximum average shear rate within 20 μm from the surface reached |50 s^−1^ (Fig. S1c). The flow is directed along +*x*, the wall lies in the *x*-*z* plane, and *y* denotes the wall-normal coordinate (Fig. 1a). The chamber height *H* is 197 μm, and we place the origin at mid-depth so that *y* ∈ [−*H*/2, *H*/2]. We define *ψ* ∈ [−π, π] as the orientation of the projection of the sperm head on the wall relative to the −*x* direction (upstream), and *θ* ∈ [−π/2, π/2] as the pitch angle describing the inclination of the head toward the wall. Flow intensity is quantified by the nondimensional ratio 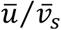, where 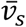is the mean swimming speed.

**Fig. 1.**
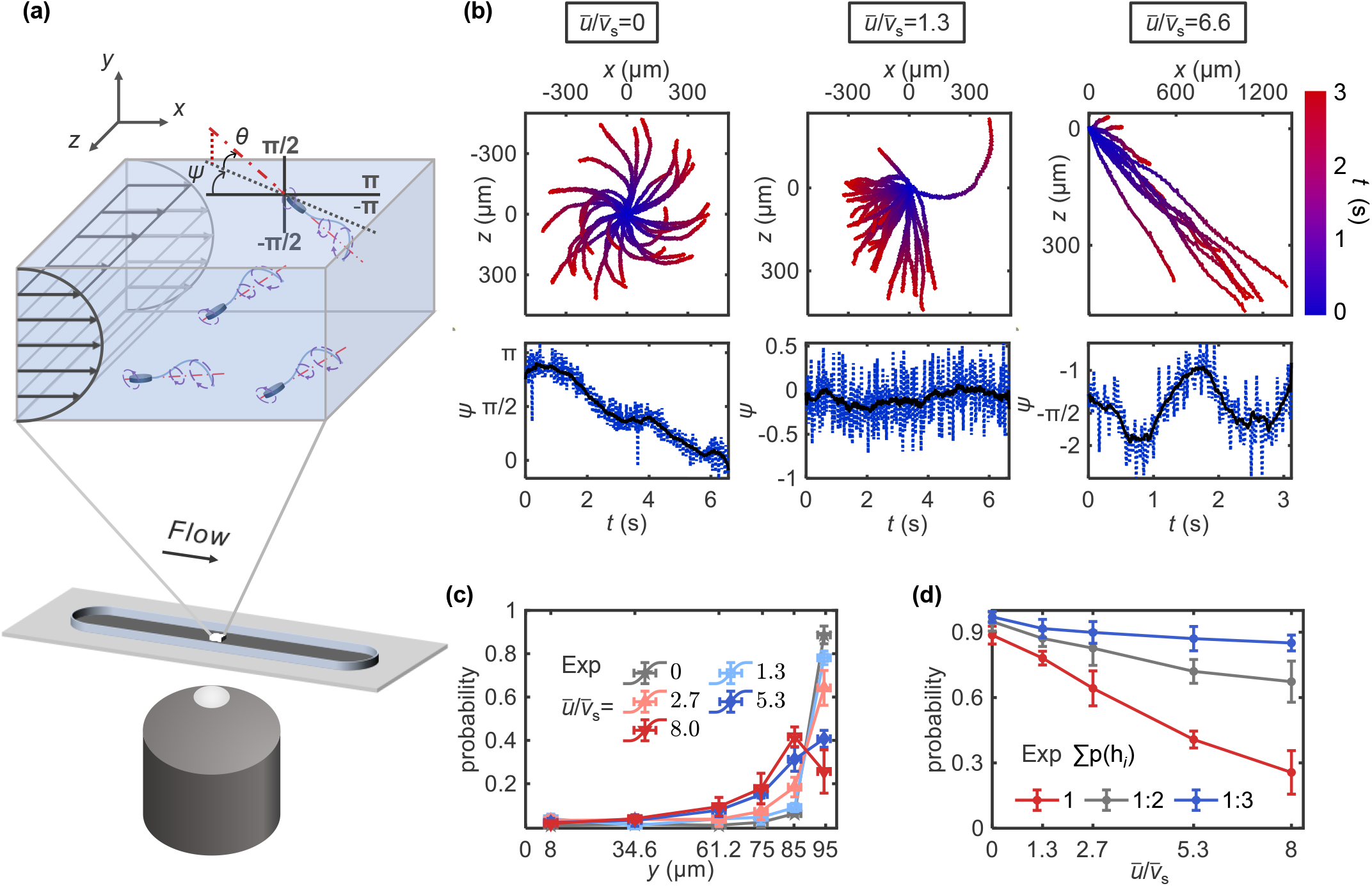
Flow-dependent sperm motility patterns and spatial distributions. (a) Schematic of the coordinate system and definition of motion parameters. (b) Typical trajectories and representative examples of distinct motility modes at three flow intensities. (c) Normalized sperm distribution measured at six wall-normal positions (*y*_h1_ = 94.5 μm, *y*_h2_ = 85.2 μm, *y*_h3_ = 74.5 μm, *y*_h4_ = 61.2 μm, *y*_h5_ = 34.6 μm, *y*_h6_ = 8 μm) for different flow intensities. (d) Cumulative near-surface probability as a function of flow intensity. Errors denote SEM.

Under near-water viscosity, most sperm exhibit rolling motion with three-dimensional flagellar beating and head rolling [40]; we therefore focus on rolling sperm throughout. Representative trajectories reveal three robust motility regimes as the flow increases (Fig. 1b). With no flow (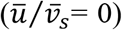), sperm swim in near-wall circles with no net transport (circular mode; Movie S1), and *ψ* varies periodically across [−π, π]. In weak flow 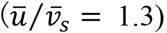, sperm align upstream and exhibit rheotaxis (|*ψ*| < π/2; Movie S2), consistent with previous reports [11,12]. At higher flow 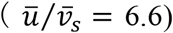, trajectories are advected downstream, yet *ψ* displays large-amplitude oscillations about −π/2. We identify this previously unreported behavior as a near-surface oscillation (NSO) mode. In NSO, the phase-contrast sharpness of the head varies periodically, indicating concomitant oscillations of the head–wall distance (Movie S3 and Fig. S2). By contrast, in the circular and rheotactic modes the head remains close to the wall.

We next quantified orientation and transport by analyzing trajectories longer than 2 s (distributions in Fig. S3). In the absence of flow, isotropic motion yields ⟨*ψ*⟩≈ 0 and negligible net drift. Upon applying flow, sperm enter a rheotactic regime. As 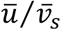 increases, alignment strengthens and then weakens: |*ψ*| initially decreases, reaches a minimum near 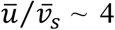, and then increases at higher flow (Fig. S3a). The drift velocities reflect the combined effects of flow advection and active swimming (Fig. S3c). Transverse drift in *z* (perpendicular to the flow) varies nonmonotonically in a manner consistent with its dependence on 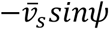. In the streamwise direction, weak flows stabilize rheotaxis and can produce net upstream migration when 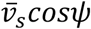exceeds ū . Consequently, a net upstream drift occurs under weak flow, as sperm actively swim against and exceed the flow velocity in the *x*-direction. As the flow increases further, rheotactic alignment deteriorates (increasing |*ψ*|), and advection dominates, leading to net downstream transport.

### Persistence of Surface Accumulation at High Shear

—Beyond rheotaxis, many microswimmers exhibit surface accumulation, typically quantified by the cell number density profile [18-20]. The emergence of the NSO mode at high flow raises a critical question: does strong shear disperse sperm into the bulk, or can near-wall accumulation persist? We measured sperm density at six discrete distances from the wall using a 20× objective (depth of field 3.0 μm) while systematically varying the flow intensity (Fig. 1c). Densities were averaged over five independent experiments and normalized to yield the spatial distribution.

In the absence of flow, sperm show pronounced near-wall enrichment, consistent with strong boundary interactions reported previously [18,21]. As flow increases, the distribution broadens and the near-wall peak is reduced, indicating partial dispersion into the bulk. Nevertheless, across all flow intensities, near-surface densities remain consistently higher than those in the bulk. Interestingly, at the highest flows, a distinct peak emerges in the distribution.

To quantify flow effects on sperm spatial distribution, we computed the cumulative probability of finding sperm within a near-surface layer (Fig. 1d). This probability decreases with increasing flow intensity, but the magnitude of the drop depends strongly on layer thickness: integrating over more near-wall bins substantially attenuates the decline. In particular, the cumulative probability for the three innermost bins remains nearly constant even at the highest shear rates. These results demonstrate that rolling sperm maintain robust near-wall occupancy under strong flows, potentially mitigating washout and preserving surface-guided navigation.

### Modeling of Sperm in Flow Fields

To elucidate the physical mechanisms underlying the observed motility transitions, we constructed a minimal orientation model incorporating (i) near-wall hydrodynamic interactions, (ii) shear-driven Jeffery rotation, and (iii) steric flagellar–wall collisions. The sperm head is modeled as a rigid prolate spheroid with semi-major axis *a* = 5 μm and semi-minor axis *b* = 1.5 μm (Fig. S4a). Propulsion is applied at an effective propulsive center, *C*, a fixed distance *d*_c_ = 13.5 μm from the head center *C*_h_. The flagellar beat is represented by two feature points, *F*_1_ and *F*_2_, located at axial distances *d*_1_ = 10 μm and *d*_2_ = 30 μm from *C*. These points oscillate transversely out of phase with amplitude *A* = 11 μm and frequency *f* = 21 Hz [41]. The swimmer orientation is described by the in-plane angle *ψ* (head projection relative to upstream) and the pitch angle *θ* (inclination toward the wall). Their dynamics are written as a superposition of hydrodynamic (*H*) Jeffery (*J*), and collision (*C*) contributions,

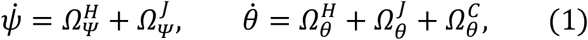

where *Ω* denotes characteristic angular speeds.

The near-wall hydrodynamic terms (Fig. S4b) are modeled by the leading far-field singularities generated by the swimmer. A force-dipole contribution (strength *α*) captures wall attraction and relaxation toward a preferred pitch angle *θ*_0_ [12,42], while a rotlet-dipole contribution (strength *β*) produces the circular trajectories observed in quiescent conditions [12,23,40]. Using image-system scaling with distance to the walls, we write

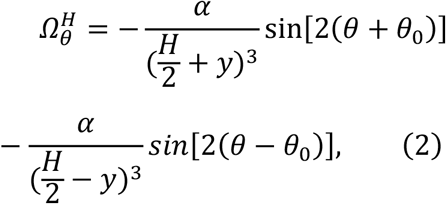

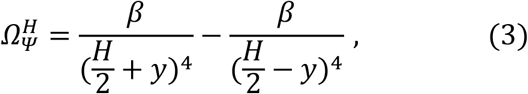

where *y* is measured from the chamber mid-plane and the two terms account for interactions with the upper and lower walls.

The flow-induced contribution (Fig. S4c) is described by Jeffery rotation of a spheroid in shear flow with local shear rate 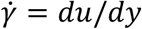[29,43]:

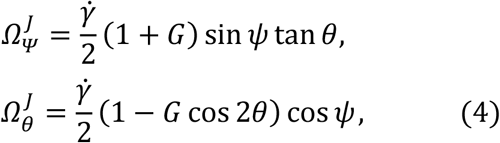

where *G* is the spheroidal shape factor.

Finally, we incorporate steric flagellar–wall collisions (Fig. S4d). When a flagellar node *F*_*i*_ enters a near-wall interaction layer of thickness *δ*_0_ (*i, e*., ||*y*_*i*=1,2_| − *H*/2| ≤ *δ*_0_) and its instantaneous normal velocity *v*_*i*_ = *dy*_*i*_/*dt* is directed toward the nearest wall, we apply an inelastic collision rule that generates a wall-directed reorientation torque. At the level of orientation dynamics, this is captured by

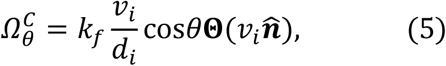

where 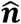is the inward wall-normal direction, **Θ** is the Heaviside function enforcing “approach-only” collisions, and *k*_*f*_ sets the effective collision inelasticity. This steric term provides a repulsive reorientation mechanism that competes with hydrodynamic attraction 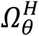and shear-driven rotation 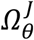, and is crucial for reproducing the persistence of near-surface swimming at high shear.

### Simulations Reproduce Sperm Motion Characteristics Observed in Experiments

—We performed 3D Langevin simulations by integrating the hydrodynamic, steric, and shear-induced torques with rotational diffusion *D*_*r*_ (see Materials and Methods). Over the experimental flow range (Fig. S3), simulations reproduce the key trends in Fig. 2: the nonmonotonic alignment response (peak shift in the *ψ* distribution; Fig. 2a), drift velocities along *x* and *z* (Fig. 2b), and the redistribution of swimmers toward the chamber center, including the emergence of a secondary peak at high flow (Fig. 2c). The near-surface occupancy, quantified by the summed probability in near-wall bins, also generally agrees with the data (Fig. 2d). Collectively, these agreements show that the minimal model captures the dominant physics across flow regimes.

**Fig. 2.**
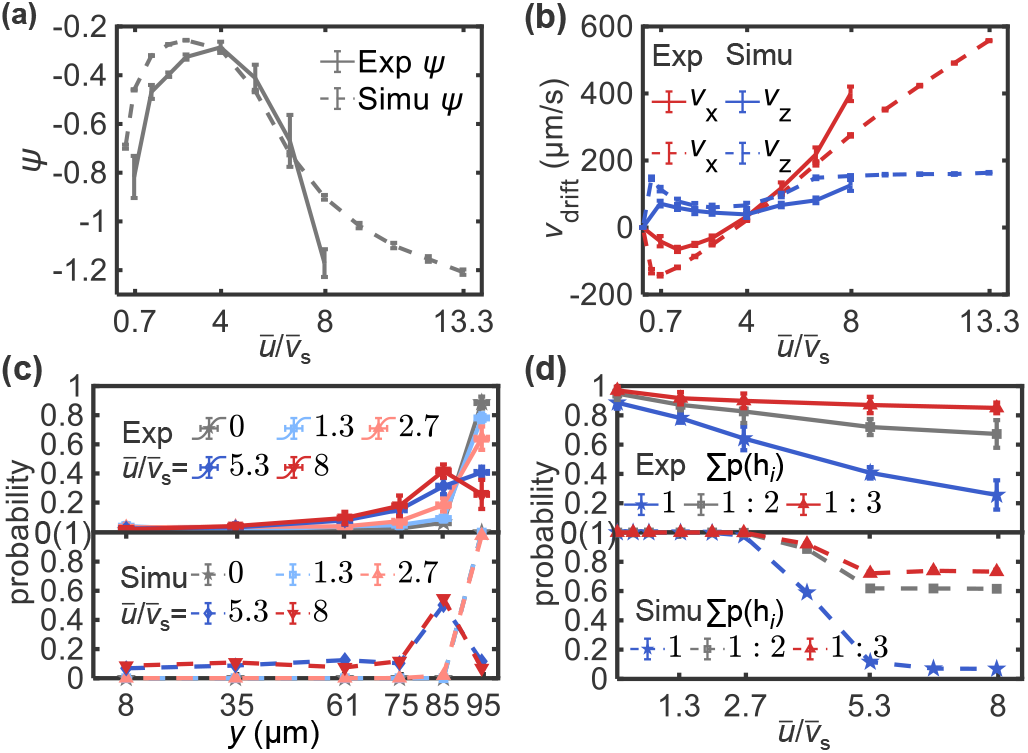
Comparison of experimental and simulation results. (a) Characteristic angle *ψ* as a function of flow intensity. (b) Drift velocity in the *x-* and *z-*directions as a function of flow intensity. (c) Normalized sperm distribution at six wall-normal positions across various flow intensities. (d) Cumulative near-surface probability as a function of flow intensity. Errors denote SEM.

To link these population statistics to single-cell dynamics, we classified simulated trajectories into six modes based on the time evolution of the wall-normal position *y* and the orientation angles *θ* and *ψ*. Representative trajectories near the top wall are shown in Fig. 3a&b. At weak or zero flow, cells exhibit a circular swimming state (CSS; Type I), with a stable wall-directed pitch (*θ* > 0). With increasing shear they enter a rheotaxis state (RS; Type II), where *ψ* fluctuates weakly about a negative mean with |*ψ*| < π/2 while maintaining an approximately constant mean pitch *θ*_0_ similar to Type I. In this regime, setting 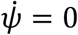 in Eq. 1 yields:

**Fig. 3.**
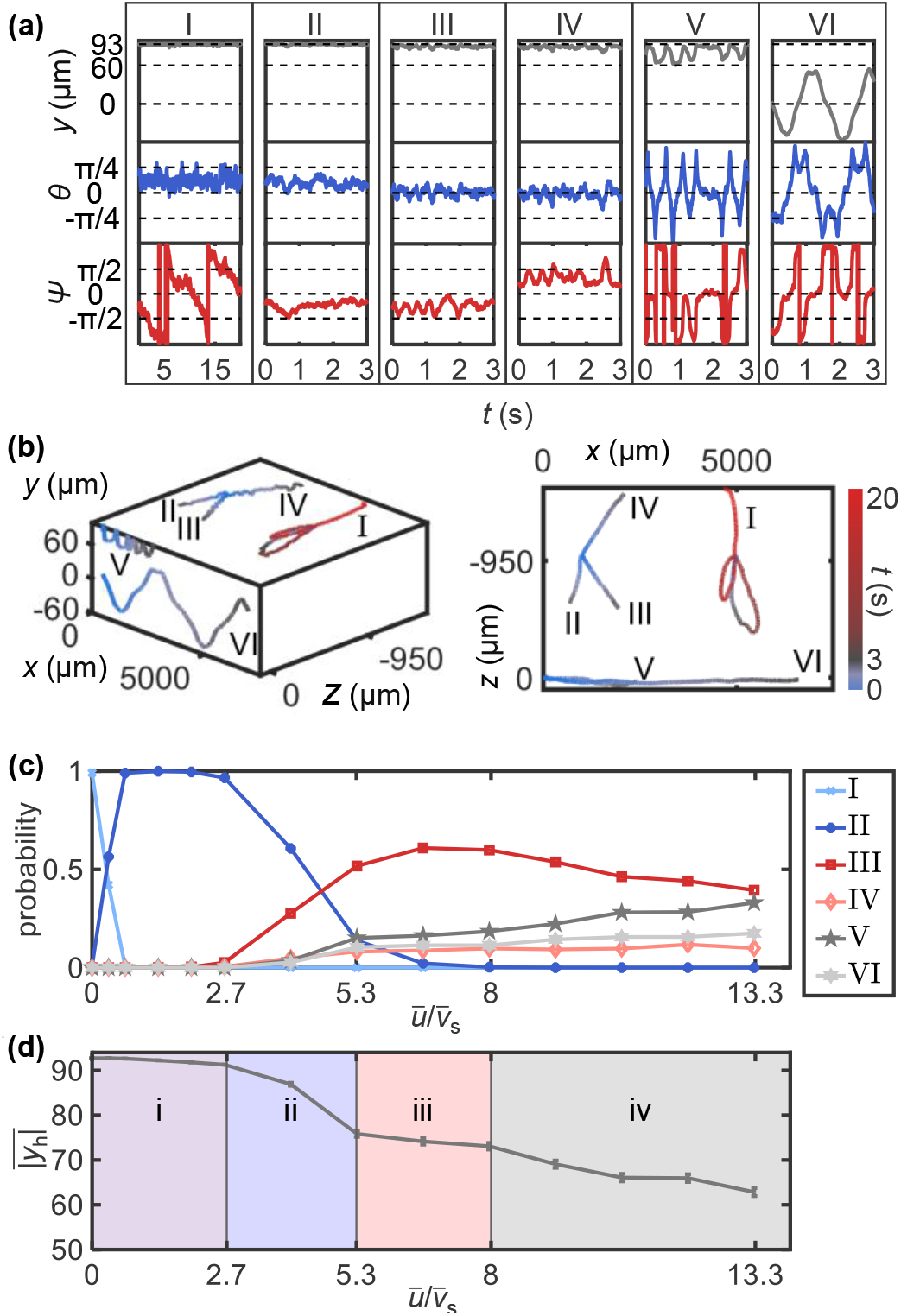
Classification and statistics of simulated sperm motility modes. (a) Definitions of the six motion patterns: (I) Circular swimming state (CSS), in the absence of flow; (II) Rheotaxis state (RS), at low flow intensity; (III) Leftward near-surface oscillation (LNSO), at high flow intensity; (IV) Rightward near-surface oscillation (RNSO), at high flow intensity; (V) Off-centerline bulk flow oscillation (OC-BFO), at very high flow intensity; Centerline-crossing bulk flow oscillation (CC-BFO), at very high flow intensity. (b) 3D trajectories for the six modes in (a), along with projections onto the *x*–*z* plane. (c Probability of each mode as a function of flow intensity. (d) Mean simulated absolute *y*-coordinate (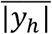) as a function of flow intensity.

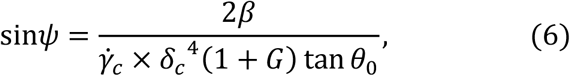

where *δ*_*c*_ and 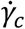 are the wall distance and shear rate evaluated at the propulsive center *C*. For weak flows both parameters vary weakly, giving 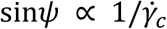, consistent with 2D cases [12].

At intermediate shear, the rheotactic fixed point destabilizes as swimmers move slightly away from the wall and the dipolar interactions weaken. The dynamics then exhibit near-surface oscillations (NSO), in which *θ* repeatedly changes sign rather than remaining locked at *θ* > 0. We distinguish leftward NSO (Type III; *ψ* < 0) and rightward NSO (Type IV; *ψ* > 0). Type III dominates at lower shear, while Type IV occurrence increases with flow intensity. At higher shear, swimmers are far enough from the wall that wall-mediated terms are negligible and trajectories become bulk flow oscillations (BFO; Types V/VI), characterized by large-amplitude oscillations of *ψ* and *θ*. We further separate BFO into off-centerline (Type V) and centerline-crossing (Type VI) modes, consistent with previous simulations of elongated microswimmers in flow [24,44].

Competition among these modes produces a nontrivial dependence of the absolute *y*-coordinate 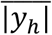on flow intensity (Fig. 3d). We identify four stages: (i) a weak decrease in 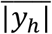as NSO first appears; (ii) a rapid drop when BFO becomes accessible; (iii) a buffer zone where 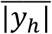plateaus despite increasing flow 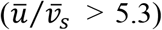, where mode fractions remain approximately constant (Fig. 3c) and the behavior is robust to parameter variations (Fig. S5); and (iv) renewed dispersal at the highest flows as bulk oscillations increase. This buffer zone provides a mechanism for maintaining near-surface proximity—and thus potential guidance by surface cues—over a finite range of elevated shear.

### Discussion

—Navigation in dynamic fluid environments is a central problem in active matter physics. For mammalian sperm, rheotaxis provides a robust long-range cue, whereas chemical and thermal gradients are often weak or unstable in vivo [1-3]. Most existing studies, however, focus on low-flow conditions [36-39]. By exploring physiologically relevant high-shear conditions, we have uncovered a rich dynamical landscape defined by the competition between hydrodynamic capture and flow-induced instability.

Our central finding is the identification of the Near-Surface Oscillation (NSO) state, a distinct dynamical mode that acts as a bridge between stable rheotaxis and bulk oscillation. Unlike bacteria or simple active colloids, which are typically dispersed from boundaries by strong flows [24,27], sperm exhibit a remarkable ability to maintain wall proximity even at high shear rates. This persistence is mechanistically underpinned by the hydrodynamic buffer zone—a regime where the destabilizing effects of Jeffery orbits are counterbalanced by steric collisions and residual dipolar attraction. This mechanism effectively caps vertical drift, creating a metastable state that prevents rapid washout.

The nonmonotonic response of sperm alignment to flow intensity highlights the complexity of this interaction. The initial enhancement of alignment followed by the onset of oscillatory instabilities suggests that the sperm flagellum has evolved to exploit, rather than merely resist, hydrodynamic shear. This buffer zone likely represents a critical evolutionary adaptation, ensuring that sperm remain within the boundary layer where flow velocities are lower and surface guidance cues are available, thereby maximizing the probability of fertilization.

From a theoretical perspective, our minimal model demonstrates that these complex behaviors emerge from the superposition of three fundamental physical components: dipolar hydrodynamics, shear-induced rotation, and steric repulsion. Quantitative agreement between experiments and simulations implies that robust surface retention and observed mode transitions do not require explicit sensory feedback; they are mechanical consequences of swimmer shape and flagellar beat-induced forcing.

These insights have broad implications beyond reproductive biology. Understanding how shape and beat kinematics confer resistance to shear washout offers design principles for artificial microswimmers intended for drug delivery in vascular environments.

Furthermore, the characterization of the high-shear buffer zone provides a physical basis for optimizing microfluidic sperm sorting technologies [45-48], moving beyond simple rheotactic selection to regimes that test the dynamic stability of the swimmer.

This work was supported by Fundamental and Interdisciplinary Disciplines Breakthrough Plan of the Ministry of Education of China (JYB2025XDXM502), National Natural Science Foundation of China (12474204), USTC Research Funds of the Double First-Class Initiative (YD2030002501), and Guizhou Provincial Major Scientific and Technological Program XKBF (2025)010.

