## Supplemental Material for "Shear-Induced Oscillations and Hydrodynamic Buffering Stabilize Sperm Surface Navigation"

### Materials and methods

**Sperm Sample Preparation.** A standard swim-up method was used to obtain sperm with good motility [1]. A 200 μL aliquot of frozen semen from a Simmental bull (purchased from a native bull station) was first thawed at 37°C, then carefully added to the bottom of 1 mL Boviwash solution (Nidacon), which had been pre-incubated (37°C, 5% $CO_2$) for over 4 hours. The centrifuge tube containing the sperm sample and Boviwash solution was incubated at 37°C with 5% $CO_2$ for 1 hour to allow motile sperm to swim into the upper layer of the solution. The supernatant (750 μL) was then carefully aspirated and washed twice (300 rpm × 5 min) to remove large impurities and dead sperm. The resulting sperm pellet was resuspended and diluted to a final surface density of ~ 5-25 × $10^{-6}/\mu m^2$.

**Microfluidic Channel Construction**. The microfluidic channel is mainly formed by a drilled glass slide, double-sided tape, and a coverslip. Two holes were manually drilled into the glass slide to serve as the channel inlet and outlet, which were connected to microfluidic tubes sealed with instant adhesive (3M Scotch-Weld). The glass slide and coverslip were combined using a custom-cut double-sided tape as a spacer, forming an inner channel with dimensions of 5 cm length × 5 mm width × 197 μm height. The outer edges of the tape were further sealed with vacuum grease. The sperm sample and solution were introduced into the channel using a microfluidic pump, which also regulated the flow field within the channel.

**Individual Sperm Trajectories Acquisition and Processing.** We used a Nikon Ti2 microscope to observe sperm near-surface motion at 10× magnification. Sperm movement was recorded at 60 fps under phase-contrast illumination with a high-frame-rate camera, allowing for an acquisition time of 10 s over the full field of view (2000 × 2000 μm). Sperm head trajectories were extracted from the recorded videos using a custom-developed Matlab program. All experiments recording sperm motion were performed at 37°C using a custom-designed constant temperature device [2].

**Calibration of Flow Field.** The flow rates ($q$) in the microfluidic channel were adjusted to four discrete values: 16, 32, 64, and 96 μL/min. The liquid source was Boviwash solution mixed with 1 μm latex beads (Polysciences) for flow visualization. The focal plane of the objective was positioned at 13.3, 24.0, 37.3, 63.9, and 90.5 μm away from the upper surface, respectively. At each focal depth, videos containing more than 1000 frames were recorded. We developed a Matlab program to track the beads in the flow field, and obtain the mean flow velocity at each position (Fig. S1a). The observation area was chosen far from the sidewalls, allowing the flow field to be approximated as a planar Poiseuille flow:

$$u = \left(\frac{4u_{max}}{H^2}\right)\left(\frac{H}{2} - y\right)\left(\frac{H}{2} + y\right), \tag{1}$$

where $u_{max}$ denotes the maximum laminar flow velocity. The mean flow velocity in the $xy$-cross section can be calculated as:

$$\bar{u} = \frac{1}{H}\int_{-\frac{H}{2}}^{\frac{H}{2}} u\, dy = \frac{2u_{max}}{3}. \tag{2}$$

After obtaining $u_{max}$ by fitting the flow velocity data to Eq. 1 at various flow rates, we found a nearly linear relationship between $\bar{u}$ and $q$ (Fig. S1b), which can be fitted by $\bar{u} = kq$. The fitted value of $k$ is ~17.7. Consequently, the shear rate can be calculated as:

$$\dot{\gamma} = \frac{du}{dy} = \frac{d\left[\left(\frac{6\bar{u}}{H^2}\right)\left(\frac{H}{2} - y\right)\left(\frac{H}{2} + y\right)\right]}{dy} = -\frac{12kqy}{H^2}. \tag{3}$$

The relationship between the mean absolute shear rate ($\dot{\gamma_{20}}$) in Poiseuille flow at 20 μm from the surface and the pumping rate ($q$) is given by: $\overline{\dot{\gamma_{20}}} = u_{y=(\frac{H}{2}-20)}/20 = 0.48q$ (Fig. S1c). Thus, under the experimental pumping rates, the maximum shear rate can reach 46.1 $s^{-1}$ at 96 μL/min.

**Sperm Distribution Measurement in Flow Fields.** To ensure a sufficient signal-to-noise ratio and keep image blur due to sperm net drift below 1 μm in each frame, we adjusted the frame rate and exposure time according to the flow intensity (Table S1). At six individual positions ($y_{h1}$ = 94.5 μm, $y_{h2}$ = 85.2μm, $y_{h3}$ = 74.5 μm, $y_{h4}$ = 61.2 μm, $y_{h5}$ = 34.6 μm, $y_{h6}$ = 8 μm), movies were recorded for 1 s each at 20× magnification to identify overlapping sperm in two-dimensional images. Sperm within the objective's depth of field (3.0 μm) were detected using a custom-developed MATLAB program based on grayscale analysis. The detection results were compared with the original movies in ImageJ to manually remove dead and agglomerated sperm. The number of individual motile sperm was then manually counted frame by frame. The final sperm density was calculated as the average over multiple frames. To ensure data reliability, single sperm counts (per experiment under specific flow conditions) ranged from 10 to 81, with the cumulative count across five replicates exceeding 131 (131-249) for each flow field.

**Brownian Dynamics (BD) Simulation of Sperm Motion.** We first considered the combined effects of force dipole, rotlet dipole, Jeffery orbits, and noise. For each timestep, the changes in $\psi$, $\theta$, and the coordinates of $C$ can be described as follows:

$$d\psi = \psi(t+dt) - \psi(t) = \left(\Omega_\psi^H + \Omega_\psi^J\right)dt + \frac{\sqrt{2D_r\,dt}\eta_\psi}{\cos\theta} =$$

$$\left[\frac{\beta}{(\frac{H}{2}+y)^4} - \frac{\beta}{(\frac{H}{2}-y)^4} + \frac{\dot{\gamma}}{2}(1+G)\sin\psi\tan\theta\right]dt + \sqrt{2D_r\,dt}\eta_\psi/\cos\theta,$$

$$\mathrm{d}\theta = \theta(t+dt) - \theta(t) = (\Omega_\theta^H + \Omega_\theta^J - \tan\theta\, D_r)dt + \sqrt{2D_r\,dt}\eta_\theta$$

$$= \left\{\begin{matrix} -\left[\frac{\alpha}{(\frac{H}{2}+y)^3}\right]\sin[2(\theta+\theta_0)] - \left[\frac{\alpha}{(\frac{H}{2}-y)^3}\right]\sin[2(\theta-\theta_0)] \\ +\frac{\dot{\gamma}}{2}(1-G\cos 2\theta)\cos\psi - \tan\theta\, D_r \end{matrix}\right\} dt + \sqrt{2D_r\,dt}\eta_\theta,$$

$$dx = x(t+dt) - x(t) = (u(y) - v_s\cos\theta\cos\psi)dt,$$

$$dy = y(t+dt) - y(t) = (v_s\sin\theta)\,dt,$$

$$dz = z(t+dt) - z(t) = (-v_s\cos\theta\sin\psi)dt, \qquad (4)$$

where $\eta_\psi$ and $\eta_\theta$ are independent Gaussian white noises with unit variance. The noise term is constructed to ensure that the sperm's orientation vector remains on the unit sphere.

For sperm-wall collisions, we sorted three classes of effects by priority: Class I collisions occur when $\delta_c \le \delta_{c0}$, where $\delta_{c0}$ denotes a distance threshold. After one timestep iteration, if $\delta_c(t) > \delta_{c0}$ and $\delta_c(t+dt) \le \delta_{c0}$, steric hindrance is applied by setting

$$y(t+dt) = \mathrm{sgn}[y(t)](\frac{H}{2} - \delta_{c0}). \qquad (5)$$

After resolving Class I collisions and iterations, Class II collisions are assumed to occur when the nearest distance between the sperm head and the surface is no more than $\delta_0$, that is,

$$\frac{H}{2} - |y_h(t)| - |a^2sin^2\theta + b^2cos^2\theta| > \delta_0 \qquad (6)$$

and

$$\frac{H}{2} - |y_h(t+dt)| - |a^2sin^2\theta + b^2cos^2\theta| \le \delta_0, \qquad (7)$$

whereupon we set

$$y(t+dt) = \mathrm{sgn}[y(t)](\frac{H}{2} - \delta_0 - |d_c\sin\theta| - |a^2sin^2\theta + b^2cos^2\theta|). \quad (8)$$

Finally, after resolving Class I and II collisions and iterations, Class III collisions correspond to collisions between the flagellar points and the surface, which occur when the following conditions holds simultaneously:

$$\left| |y_{i=1,2}| - \frac{H}{2} \right| \leq \delta_0, \tag{9}$$

where $\delta_0$ is the threshold distance defining collision proximity, and

$$y_i v_i > 0, \tag{10}$$

where $v_i = dy_i/dt$ is the velocity of $F_1$ or $F_2$ in the $y$-direction, pointing toward the nearest wall. When conditions of Eqs. 9 and 10 are simultaneously met, we defined a characteristic angular velocity $\omega_i$ as:

$$\omega_{i=1,2} = k_f \cos\theta \, (\frac{\mathrm{d}y}{dt} + 2\pi f A \cos\theta \cos(2\pi f t + i\pi))/d_i. \tag{11}$$

For each timestep, if there exists a flagella-wall collision, the changes in $\theta$ and the coordinates of $C$ can be described as follows (adapted from Eq. 4):

$$\mathrm{d}\theta = (\Omega_\theta^H + \Omega_\theta^J - \tan\theta \, D_r)dt + \sqrt{2D_r\, dt}\eta_\theta$$

$$= \left\{ \begin{matrix} -\left[\dfrac{\alpha}{(\frac{H}{2}+y)^3}\right]\sin[2(\theta+\theta_0)] - \left[\dfrac{\alpha}{(\frac{H}{2}-y)^3}\right]\sin[2(\theta-\theta_0)] \\ +\dfrac{\dot{\gamma}}{2}(1 - G\cos 2\theta)\cos\psi - \tan\theta\, D_r \end{matrix} \right\} dt$$

$$+\sqrt{2D_r\, dt}\eta_\theta + \omega_i dt,$$

$$dy = (v_s \sin\theta)\, dt + \omega_i dt d_i \cos\theta,$$

$$dx = (u(y) - v_s \cos\theta \cos\psi)dt - \omega_i dt d_i \mathrm{sgn}(y) |\sin\theta| \cos\psi,$$

$$dz = (-v_s \cos\theta \sin\psi)dt - \omega_i dt d_i \mathrm{sgn}(y) |\sin\theta| \sin\psi, \tag{12}$$

where $i = 1$ and 2 correspond to the two feature points on the flagellar projection.

The simulation domain is defined as $x \in [0, L]$, $y \in [-H/2, H/2]$, $z \in [0, L]$, where $L$ = 50000 μm and $H$ = 197 μm. In the $x$ and $z$-directions, periodic boundary conditions are applied to avoid data overload, so sperm motion is not restricted by thresholds in these directions. Initial conditions are set as follows: $x|_{t=0} \sim$ U(0, $L$), $y|_{t=0} \sim$ U($-H/2 + 20$, $H/2 - 20$), $z|_{t=0} \sim$ U(0, $L$), $\psi|_{t=0} \sim$ U(-π, π), and $\theta|_{t=0} \sim$ U(-π/2, π/2), where U($a$, $b$) denotes a uniform distribution on [$a$, $b$].

Parameter values used in the simulation are summarized in Table S2. The force dipole factor is given by $\alpha = 3p/128\pi\eta$, where $p$, the dipole strength, is estimated to be 280 pN·μm [3]. The viscosity $\eta$ of the experimental solution is approximated as that of water, about 0.0007 Pa·s at 37°C, yielding $\alpha \approx 2984$ μm$^3$s$^{-1}$. The stable average pitch angle $\theta_0$, resulting from the conical envelope of the flagellar wave, was measured previously to be between 10° and 20° [4]. Here, we used $\theta_0 \sim 20°$. The distance threshold $\delta_0$ is estimated to be 2.5 μm, considering the sperm size. The distances $d_c$, $d_1$ and $d_2$ are estimated to be 13.5, 10 and 30 μm, respectively, as described in the modeling section. Consequently, the distance threshold $\delta_{c0}$ can be calculated as $d_c\sin\theta_0 + |a^2\sin^2\theta_0 + b^2\cos^2\theta_0| + \delta_0$, which is ~9.33 μm. The amplitude $A$ of two feature points $F_1$ and $F_2$ on the flagellum is estimated to be 11 μm, and the frequency $f$ is ~21 Hz [2]. To match the experimental results, the flow field parameter $G$ and the correction factor for inelastic collision $k_f$ are manually set to 0.75 and 0.16, respectively.

To estimate the rotational diffusion coefficient $D_r$, we tracked sperm trajectories near the surface in the absence of flow. We quantified $D_r$ by computing the mean-squared angular displacement (MSAD) from tracking data (Fig. S6a), which can be fitted by (Fig. S6b):

$$\mathrm{MSAD} = \langle[\psi(t)] - \psi(0)]^2\rangle = \omega^2 t^2 + 2D_r t, \qquad (13)$$

where $\omega$ denotes the angular velocity of sperm due to the rotlet dipole effect, with a fitted value of ~0.58 rad/s, and the fitted value of $D_r$ is ~0.16 s$^{-1}$. The rotlet dipole effect coefficient $\beta$ can be calculated using $\omega\delta_{c0}^4$, which is ~4395 μm$^4$ s$^{-1}$. The sperm trajectories (31 sperm) were further used to obtain the average swimming speed of sperm ($\bar{v}_s$), which is ~213.3 μm s$^{-1}$. The timestep of simulation was set to 0.001 s, much shorter than the rotational diffusion timescale $1/(2D_r) = 3.125$ s. Every 20 seconds, the time evolution (from time zero) of the average $y$-coordinates (absolute value) of 1000 simulated sperm trajectories was displayed to determine whether the sperm distribution had reached equilibrium. Once equilibrium was established, the simulation was terminated, and the sperm distribution during the last 20 seconds was averaged and then compared with experimental results.

**Classification of Sperm Motility States.** To classify sperm trajectories into six motility states under varying flow conditions, we established five quantitative criteria based on orientation, hydrodynamic interaction, and spatial dynamics. Classification proceeded in two ordered stages: full-trajectory evaluation (20 s) followed by segment-based analysis (period length: 1 s). Parameters of classification are:

1) Backtracking probability, the probability that the absolute orientation angle exceeds π/2 ($P(|\psi|>\pi/2)$) during the 20 s trajectory, serves as a criterion for distinguishing between circular and directional motion patterns.

2) Wall-directed orientation ratio (WOR), defined as the fraction of the trajectory when the sperm head points toward the surface.

3) Hydrodynamic interaction coefficient (HIC), which reflects surface-induced hydrodynamic interactions and is defined as:

$$\mathrm{HIC} = \begin{cases} \dfrac{\alpha}{(\frac{H}{2}+y)^3} (y < 0) \\ \dfrac{\alpha}{(\frac{H}{2}-y)^3} (y > 0) \end{cases} \tag{14}$$

Under no-flow conditions, HIC is approximately 2.36 $s^{-1}$; thus, we suppose effective surface interaction occurs if the mean HIC of a trajectory is larger than 0.47 (20% of the reference value).

State classification was then conducted using the following criteria:

<table>
<tr><th>State</th><th>Full-trajectory evaluation (20 s)</th><th>Segment-based evaluation (1 s)</th></tr>
<tr><td>CSS (I)</td><td>P(|ψ|>π/2) > 0.1,<br>WOR > 80%,<br>HIC > 0.47</td><td>——</td></tr>
<tr><td>RS (II)</td><td rowspan="6">Others</td><td>WOR > 80%,<br>HIC > 0.47</td></tr>
<tr><td>LNSO (III)</td><td>WOR ≤ 80%,<br>HIC > 0.47,<br>P(ψ < 0) > 50%</td></tr>
<tr><td>RNSO (IV)</td><td>WOR ≤ 80%,<br>HIC > 0.47,<br>P(ψ > 0) > 50%</td></tr>
<tr><td>OC-BFO (V)</td><td>HIC < 0.47,<br>P($y_h$ < 0) = 100% or<br>P($y_h$ > 0) = 100%</td></tr>
<tr><td>CC-BFO (VI)</td><td>HIC < 0.47,<br>0 < P($y_h$ < 0) < 100%</td></tr>
</table>

We first assessed whether a complete 20-s trajectory could be classified as the CSS state. If so, it would contribute 20 counts to the statistical results. Otherwise, segment-based evaluation was applied to judge the state of each 1-s segment (contributing one count to the statistical results) of the 20-s trajectory. Through this method, more than 99% of the trajectory segments were successfully classified as one of the six motility states. There were ~20,000 segments of trajectory involved in the statistics.

**Supplementary Note 1: The Effect of Parameter Settings on Sperm Distribution under Various Flow Fields.** In the main text, we reproduced experimental results by setting appropriate parameter values in the model. Here, we investigated how variations in these parameter values affect sperm distribution in flow fields. To address this, we focused on the average ($\overline{|y_h|}$) of the sperm population in relation to different parameter values ($\overline{|y_h|}$ = 0 corresponds to the central layer of liquid, and $\overline{|y_h|}$ = $H/2$ corresponds to the surface plane).

As shown in Fig. 3d of the main text, we found that the "buffer zone" robustly exists regardless of variations in key parameters within reasonable ranges (Fig. S5). However, the relationship between $\overline{|y_h|}$ and $\bar{u}/\bar{v}_s$ is influenced by parameter settings. Typically, with smaller $k_f$—i.e., when flagella-wall collision effects are weakened—sperm escape from the wall more rapidly as flow intensity increases (Fig. S5a). Moreover, with a larger $G$, sperm display stronger anisotropic hydrodynamic responses to shear flow [5,6], thereby promoting the separation between sperm and the wall (Fig. S5b). Regarding far-field hydrodynamic interactions, stronger force dipole effects lead to more stable surface residence of sperm, as expected (Fig. S5c) [7].

### Supplementary Figures

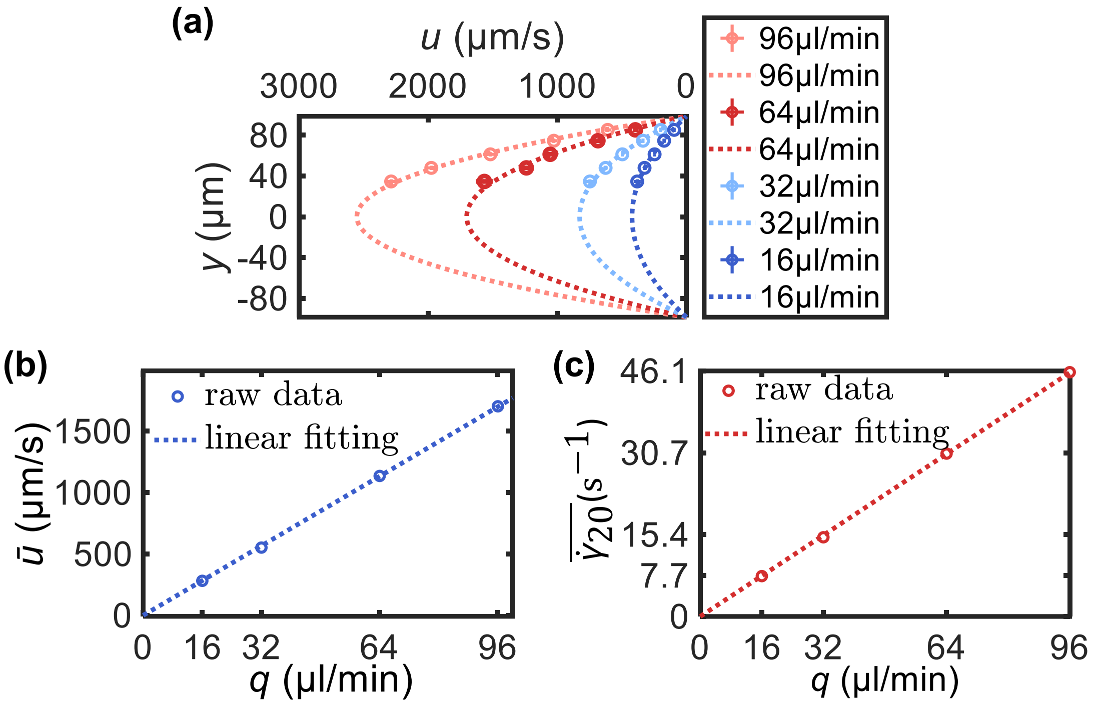


**Fig. S1. Experimental setup and flow-field calibration.** (**a**) Experimental flow velocity profiles measured at different pumping rates, overlaid with parabolic fits. (**b**) Average flow velocity versus pumping rate. The blue dotted line represents a linear fit. (**b**) Average shear rate within 20 μm from the surface versus pumping rate. The red dotted line represents a linear fit.

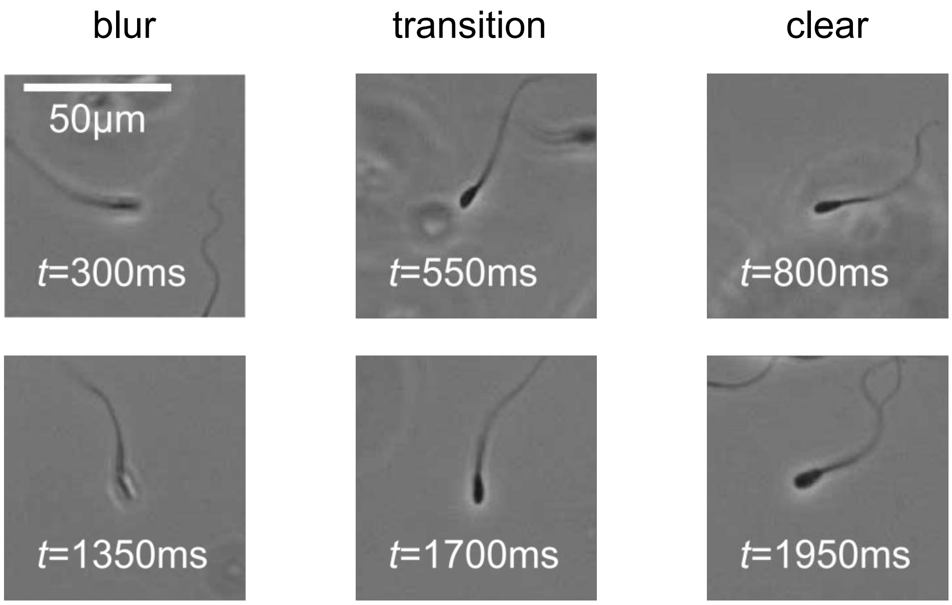


**Fig. S2. Near surface oscillation of the phase-contrast imaging clarity.** In high flow fields, sperm exhibit oscillatory motion in the *y*-direction (normal to the focal plane of the objective), resulting in periodic defocusing-refocusing cycles that generate periodic transitions between blurred and clear imaging.

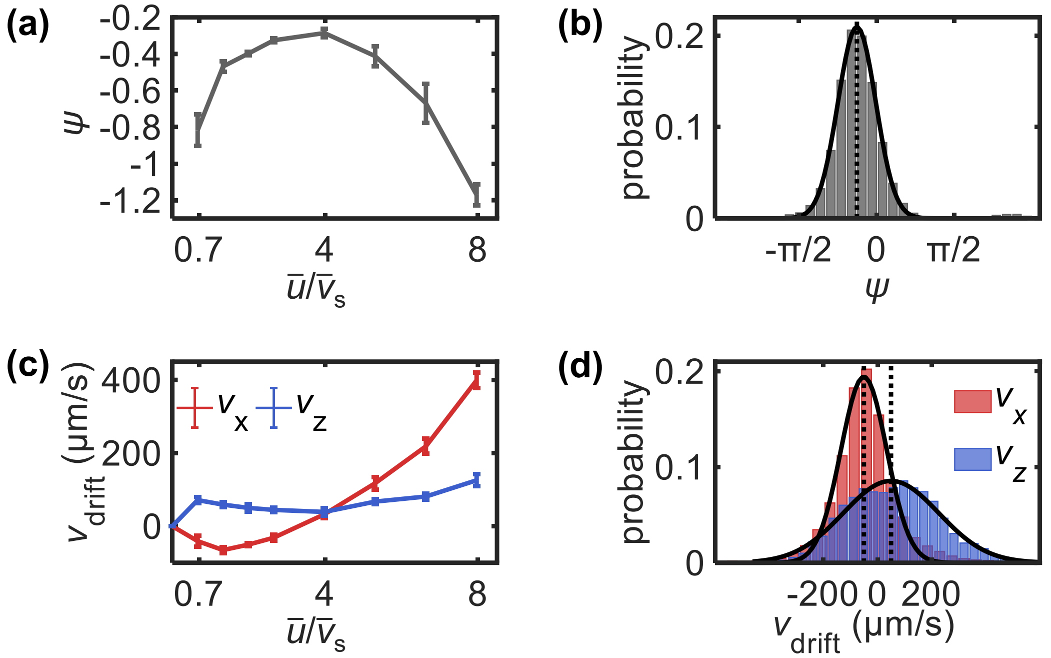


**Fig. S3. Characteristic angle $\psi$ and pure drift velocity of rolling sperm in various flow fields.** (**a**) $\psi$ in relation to flow field intensity. (**b**) Gaussian fitting of the distribution of $\psi$ at $\bar{u}/\bar{v}_s = 2.0$. The mean and standard deviation of the fits are shown in (*A*). (**b**) Pure drift velocities in the *x* and *z* directions in relation to flow field intensity. (**d**) Gaussian fitting of the distribution of drift velocities in the *x* and *z* directions at $\bar{u}/\bar{v}_s = 2.0$. The mean and standard deviation of the fits are shown in (c).

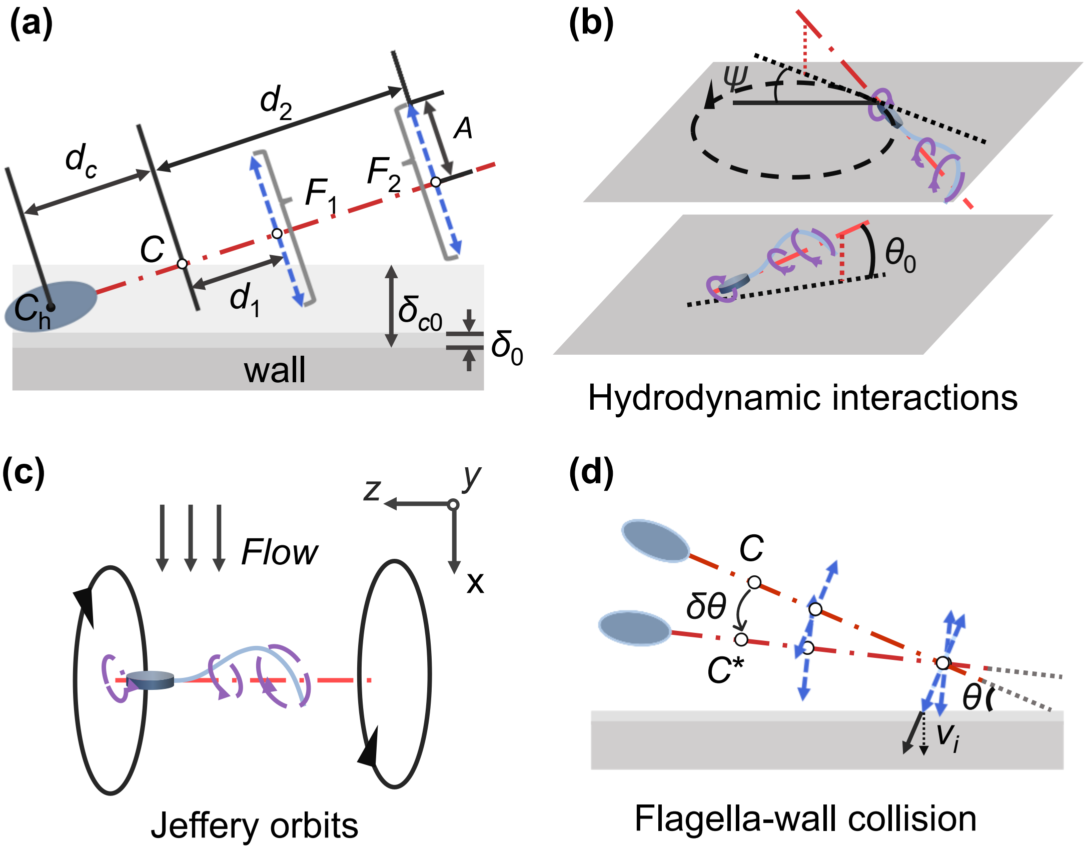


**Fig. S4. Schematic diagram of the sperm model and sperm-wall interactions in flow fields.** (**a**) Illustration of the sperm model and key parameters. (**b**) Hydrodynamic interactions between sperm and the surface. The force dipole effect induces a stable pitch angle of the sperm head, while the rotlet dipole effect causes circular motion along the surface. (**c**) Jeffery-orbit effect induced by flow fields. The direction of the periodic motion depends on the shear direction. (**d**) Flagella-wall collisions. The torque exerted by the reaction force leads to sperm reorientation toward a configuration more parallel to the surface.

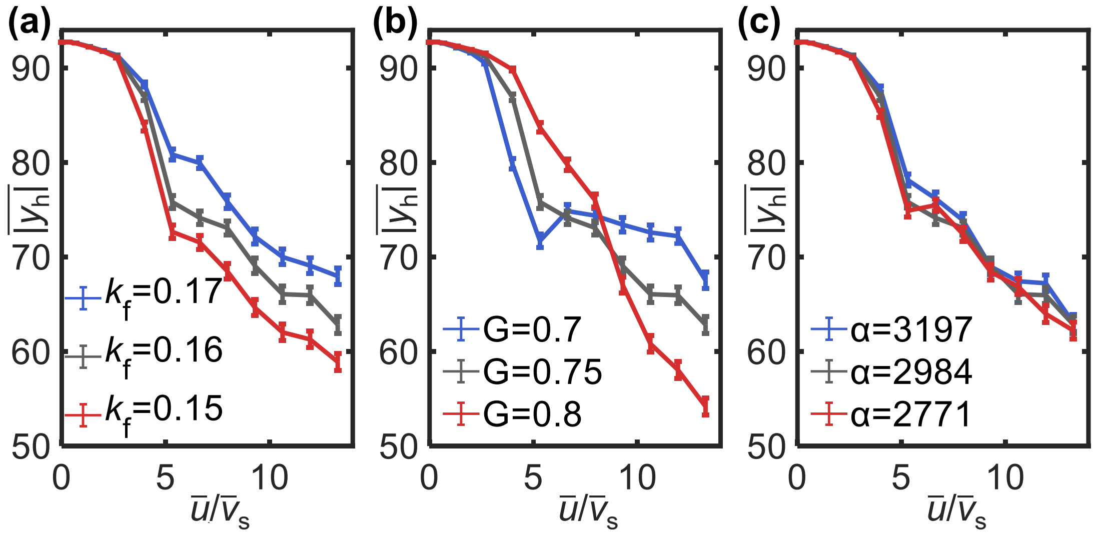


**Fig. S5. Simulated mean position (absolute value) of sperm head in relation to flow intensity with various parameter values.** (**a**) Results with varying flagellum factor $k_f$. (**b**) Results with varying flow field parameter $G$. (**c**) Results with varying force dipole factor $\alpha$.

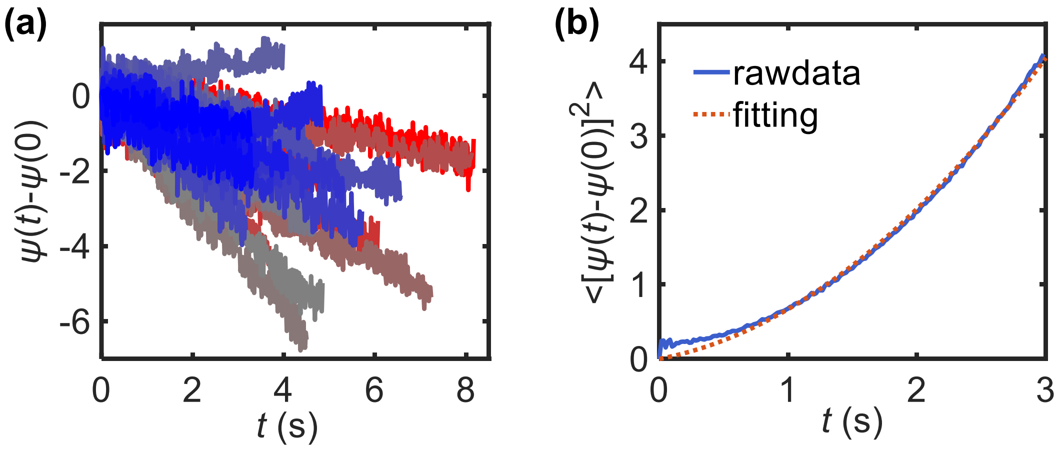


**Fig. S6. Calibration of the rotational diffusion coefficient of bovine sperm.** (**a**) Variation of $\psi$ over time ($t$) for 31 individual sperm in the absence of flow. (**b**) Mean-squared angular displacement (MSAD) versus time. The orange dotted line represents the fit according to Eq. 13 in Materials and Methods.

### Supplemental Tables

**Table S1. Frame rate and exposure time setting in relation to flow intensity for the measurement of sperm distributions.**

| Flow rate (µL/min) | Frame rate ($s^{-1}$) | Exposure time (µs) |
|---|---|---|
| 96 | 60 | 300 |
| 64 | 45 | 450 |
| 32 | 30 | 900 |
| 16 | 30 | 1800 |
| 0 | 30 | 1800 |

**Table S2. Parameter setting for simulation.**

| Parameter | Value |
|---|---|
| $\beta$ | 4395 µm$^4$ s$^{-1}$ |
| $H$ | 197 µm |
| $G$ | 0.75 |
| $D_r$ | 0.16 s$^{-1}$ |
| $\alpha$ | 2984 µm$^3$ s$^{-1}$ |
| $\theta_0$ | 20° |
| $\delta_{c0}$ | 9.33 µm |
| $\delta_0$ | 2.5 µm |
| $k_{\mathrm{f}}$ | 0.16 |
| $v_{\mathrm{s}}$ | 213.3 µm s$^{-1}$ |
| $a$ | 5 µm |
| $b$ | 1.5 µm |
| $A$ | 11µm |
| $f$ | 21 Hz |
| $d_{\mathrm{c}}$ | 13.5 µm |
| $d_1$ | 10 µm |
| $d_2$ | 30 µm |
| $dt$ | 0.001 s |

### Supplemental Movies

**Movie S1.** A typical example showing sperm circular swimming near the surface due to the rotlet dipole effect.

**Movie S2.** A typical example showing sperm upstream swimming near the surface due to the application of weak flow fields.

**Movie S3.** A typical example showing the variation in the phase-contrast imaging clarity of the sperm head due to the oscillating distance between the sperm head and the surface under high flow fields.
